# A high-throughput analysis method of microdroplet PCR coupled with fluorescence spectrophotometry

**DOI:** 10.1101/181974

**Authors:** Yanan Du, Xiao Zhao, Binan Zhao, Yan Xu, Wei Shi, Fangfang Ren, Yangyang Wu, Ruili Hu, Xiaorui Fan, Qi Zhang, Xiaoxia Zhang, Bin Shi, Huanzhen Zhao, Kai Zhao

**Author notes:** Y.N.D. and X.Z. contributed equally to this work. Correspondence: Kai Zhao.

## Abstract

Here we report a novel microdroplet PCR method combined with fluorescence spectrophotometry (MPFS), which allows for qualitative, quantitative and high -throughput detection of multiple DNA targets. In this study, each pair of primers was labeled with a specific fluorophore. Through microdroplet PCR, a target DNA was amplified and labeled with the same fluorophore. After products purification, the DNA products tagged with different fluorophores could be analyzed qualitatively by the fluorescent intensity determination. The relative fluorensence unit was also measured to construct the standard curve and to achieve quantitative analysis. In a reaction, the co -amplified products with different fluorophores could be simultaneously analyzed to achieve high -throughput detection. We used four kinds of GM maize as a model to confirm this theory. The qualitative results revealed high specificity and sensitivity of 0.5% (w / w). The quantitative results revealed that the limit of detection was 10^3^copies and with good repeatability. Moreover, reproducibility assay were further performed using four foodborne pathogenic bacteria. Consequently, the same qualitative, quantitative and high-throughput results were confirmed as the four GM maize.

## Introduction

With the increasing of the number of bimolecular samples to be analyzed, it demands a high -throughput detection method capable of both qualification and quantification. Traditional detection technology can usually only detect one target gene in one reaction. When dealing with complex nucleic acid samples, multiple reactions / detection need to be separately performed, which is time consuming and costly. Therefore, it is of great value to develop a method which enables qualitative and quantitative detection simultaneously at reasonable cost. Indeed, several multiplex detection methods of target genes had been developed and employed in the fields, such as food / feed identifications (Settanni and Corsetti 2007; Chaouachi et al. 2014; Morisset et al. 2008; Guo et al. 2011), medical diagnostics (Barken et al. 2007; Uttamchandani et al. 2009; Ge et al. 2006) and large scale sequencing (Porreca et al. 2007; Tewhey et al. 2009; Krishnakumar et al. 2008).

Although conventional multiplex PCR were always used to amplify nucleic acid samples, many problems exist against its wide use, such as preferential amplification of shorter DNA templates, interference of multiple primer pairs and limited substrates, making it incapable in quantitative and high -throughput research (Meyerhans et al. 1990; Dahl et al. 2005). Real -time qPCR has been widely used to quantify target genes with high sensitivity, specificity and a wide dynamic range (Heid et al. 1996; Higuchi et al. 1993; Wittwer et al. 1997; Holland et al.1991), the throughput in such a scheme is low because of the limited number of channels in the real -time system. DNA microarray is an approach to analyze complex nucleic samples with high throughput, the complicated procedure and expensive consumption, however, limit its feasible availability for most laboratories. Poor linearity is another weakness related with DNA microarray which limits its use in quantitative analysis (Hessner et al. 2003). Therefore, a novel qualitative, quantitative and high-throughput method for detecting multiple biological genes is on demand.

The multiplex emulsion PCR has been developed for high -throughput simultaneous amplification of several DNA targets, either used alone (Chaouachi et al. 2008; Williams et al. 2006) or combined with other methods (Guo et al. 2011; Barken et al. 2007). In emulsion PCR, the different target DNA molecules are amplified in parallel in millions of compartmentalized micro -reservoirs, which alleviates the drawbacks in conventional multiplex PCR while increasing the throughput and reducing the regent and sample consumption.

In this paper, we describe a novel high -throughput method developed in our laboratory, which combines microdroplet PCR with fluorescence spectrophotometry (MPFS). The MPFS method used the primers labeled with different fluorophores, without detection interference, so that the multitarget DNAs in one sample could be amplified in a single reaction set and analyzed both qualitatively and quantitatively on Infinite M1000 PRO, after products purification. This method has been tested with four event specific GM maize and four foodborn pathogenic bacteria with satisfaction. The MPFS method provides a new approach for qualitative, quantitative and high -throughput analysis of multitarget DNAs to a broad range of biological samples.

## Results

### The Principle of MPFS

In MPFS, each of the fluorophores has its own intrinsic excitation and emission wavelength that can be identified and measured by the Infinite M1000 PRO. Each pair of primers specific to a target sequence was labeled with a specific fluorophore that does not interfere with any other fluorophores in the same reaction set for detection purpose. Through the emulsion PCR, a target DNA was amplified and labeled with the same fluorophore. After PCR reaction, the products were purified by PCR cleanup kit to remove the free primers and other disruptive substances. Finally, the DNA products tagged with different fluorophores could be used for qualitative analysis by the fluorescent intensity determination. The relative fluorescence unit (RFU) was also measured to construct the standard curve and then the quantitative analysis could be achieved. Since a variety of different target samples were simultaneously amplified in a single PCR tube, the co -amplified products with different fluorophores could be simultaneously analyzed to achieve high -throughput detection.

### BT176, GA21, NK603 and TC1507 Singleplex Assay

#### Specificity Assay and Qualitative Detection

In order to confirm the specificity of the singleplex MPFS method, a mixed DNA template solution containing eight events (BT176 maize, GA21 maize, NK603 maize, TC1507 maize, non-GM maize, soybean, and rapeseed), no template control (NTC) and a pair of labeled primers was employed. Results are shown in the Table 1. In BT176 specificity assay, only the RFUs of BT176 genome and the positive control (BT176 plasmid) were higher than the threshold and revealed positive. While others revealed negative. GA21 maize, NK603 maize and TC1507 maize showed the similar results as BT176.

**Table 1.** BT176, GA21, NK603 and TC1507 singleplex specificity assay.

| Samples | BT176(RFU) | GA21(RFU) | NK603(RFU) | TC1507(RFU) | IVR(RFU) |
| --- | --- | --- | --- | --- | --- |
| BT176 PC | 28739(+) |  |  |  |  |
| GA21 PC |  | 7957(+) |  |  |  |
| NK603 PC |  |  | 16272(+) |  |  |
| TC1507 PC |  |  |  | 27249(+) |  |
| IVR PC |  |  |  |  | 27166(+) |
| BT176 maize | 8654(+) | 5926(+) | 686(-) | 2016(-) | 27390(+) |
| GA21 maize | 1998(-) | 1593(-) | 739(-) | 2096(-) | 28234(+) |
| NK603 maize | 1567(-) | 1838(-) | 9119(+) | 2145(-) | 28019(+) |
| TC1507maize | 2088(-) | 1663(-) | 660(-) | 23345(+) | 27441(+) |
| Wild maize | 1492 (-) | 1697(-) | 633(-) | 2111(-) | 27390(+) |
| Soybean | 1801(-) | 1432(-) | 712(-) | 2761(-) | 1052(-) |
| Rapeseed | 1844(-) | 1732(-) | 923(-) | 1953(-) | 1102(-) |
| Wheat | 2272(-) | 1613 (-) | 844(-) | 1786(-) | 883(-) |
| Rice | 2139(-) | 2290(-) | 555(-) | 2388(-) | 1171(-) |
| NTC $\bar{X}$ | 1892.72 | 1625.58 | 629.13 | 2127.92 | 1186.90 |
| NTC SD | 252.08 | 189.34 | 90.95 | 282.29 | 257.30 |
| Threshold | 2901.03 | 2382.95 | 992.95 | 3257.07 | 2216.08 |

RFU: Relative fluorescence units; PC: Positive control; NTC: No template control; $\bar{X}$ : The average RFU of NTC;
SD: Standard deviation; Threshold: $\bar{X}+4SD$ ; +: Positive signal; -: Negative signal.

The specificity of maize endogenous gene was assessed on a series of template samples. The template samples include endogenous gene plasmid (positive control), BT176 maize, GA21 maize, NK603 maize, TC1507 maize, non-GM maize, NTC, soybean, rapeseed, wheat, cotton and rice. As a result (Table 1), only the RFUs of endogenous gene plasmid, BT176 maize, GA21 maize, NK603 maize, TC1507 maize and non-GM maize were higher than the threshold and showed positive signal. Others revealed negative.

Therefore, the singleplex MPFS PCR assay demonstrated high specificity for detecting GM maize and maize endogenous gene.

#### Sensitivity Assay

For testing the sensitivity of the singleplex MPFS method, four series of relative GMO content (fortified at 10%, 5%, 1%, and 0.5%) were employed as templates, respectively. Results were shown in Table 2. As shown, all RFU values of Bt176 maize were higher than the threshold and increased along with the GMO content. GA21 maize, NK603 maize and TC1507 maize showed the similar results as BT176 maize. Of notes, all four maize samples were detected positive as low as 0.5% (w/w) GMO content that is lower than EU regulations.

**Table 2.**
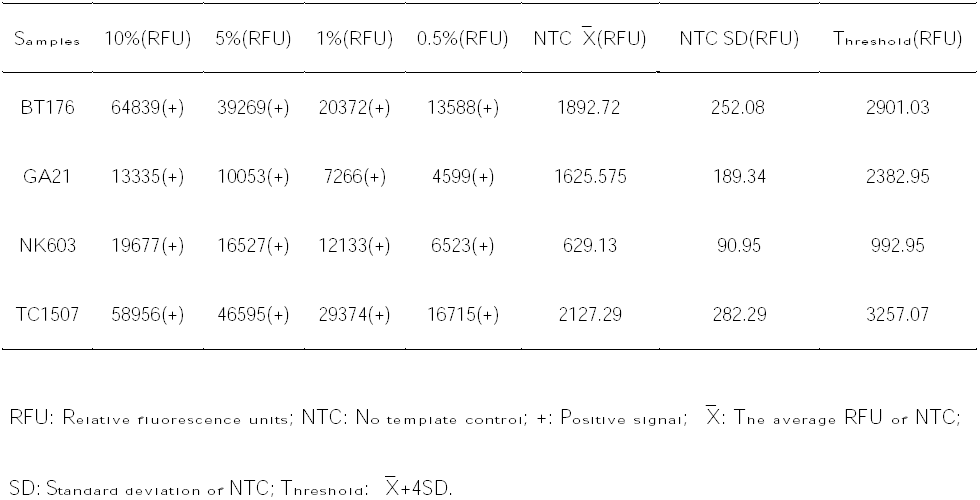
BT176, GA21, NK603 and TC1507 singleplex sensitivity assay.

| Samples | 10%(RFU) | 5%(RFU) | 1%(RFU) | 0.5%(RFU) | NTC $\bar{X}$(RFU) | NTC SD(RFU) | Threshold(RFU) |
| --- | --- | --- | --- | --- | --- | --- | --- |
| BT176 | 64839(+) | 39269(+) | 20372(+) | 13588(+) | 1892.72 | 252.08 | 2901.03 |
| GA21 | 13335(+) | 10053(+) | 7266(+) | 4599(+) | 1625.575 | 189.34 | 2382.95 |
| NK603 | 19677(+) | 16527(+) | 12133(+) | 6523(+) | 629.13 | 90.95 | 992.95 |
| TC1507 | 58956(+) | 46595(+) | 29374(+) | 16715(+) | 2127.29 | 282.29 | 3257.07 |

RFU: Relative fluorescence units; NTC: No template control; +: Positive signal; $\bar{X}$ : The average RFU of NTC; SD: Standard deviation of NTC; Threshold: $\bar{X}+4SD$ .

Therefore, the singleplex MPFS PCR assay demonstrated high sensitivity for detection of GM maize.

#### Construction of Standard Curves and Quantitative Detection

Five concentrations from 10^3^ to 10^7^copies of four GM maize and endogenous gene were used to construct singleplex standard curves with NTC as control. Results are shown in the Figure 1. In BT176 assay, the plots showed a typical log-linear standard curve and the regression correlation coefficient (R^2^) value were 0.9996, which indicated excellent relationship between the DNA template copy numbers and the RFU values. It is also true to other samples, such as GA21 maize, NK603 maize and TC1507 maize, as well as the maize endogenous gene. The limit of detection (LOD) was 10^3^copies in all the five samples. Therefore, the singleplex MPFS method demonstrated high accuracy and can be used to quantify the GM maize.

**Figure 1.**
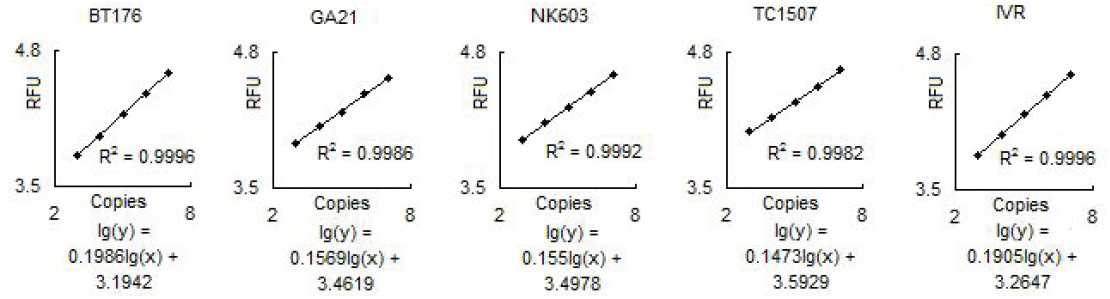
BT176, GA21, NK603 and TC1507 Singleplex standard curves. RFU: Relative fluorescence units; Copies: The copies of plasmids.

### BT176, GA21, NK603 and TC1507 Multiplex Assay

#### Specificity Assay, Qualitative Detection and High Throughput

Specificity of the 4-plex of MPFS were confirmed on the DNA materials containing four events (BT176 maize, GA21 maize, NK603 maize, TC1507 maize) in a single reaction. Results are shown in the Table 3. The RFU of four maize genomes were higher than the threshold and showed positive signals. While the NTC revealed negative. Therefore, the 4-plex MPFS assay demonstrated high throughput and the same specificity as the single MPFS assay. Overall these data showed that increasing the number of templates in the MFPS method did not affect the result of qualitative detection.

**Table 3.** BT176, GA21, NK603 and TC1507 multiplex specificity assay.

| Samples | Maize genome(RFU) | NTC $\bar{X}$(RFU) | NTC SD(RFU) | Threshold (RFU) |
| --- | --- | --- | --- | --- |
| BT176 | 20056(+) | 1166.24 | 188.22 | 1919.13 |
| GA21 | 26440(+) | 1423.03 | 310.23 | 2663.94 |
| NK603 | 28880(+) | 1004.99 | 211.64 | 1851.56 |
| TC1507 | 46763(+) | 2284.97 | 358.86 | 3720.10 |

RFU: Relative fluorescence units; NTC: No template control; $\bar{X}$ : The average RFU of NTC; SD: Standard deviation of NTC; Threshold: $\bar{X}+4SD$ ; +: Positive signal.

#### Sensitivity Assay

Sensitivity of 4-plex MPFS method was investigated on the DNA materials containing four GM maize genomes at all levels (fortified at 10%, 5%, 1%, and 0.5%) in a single tube. Results are shown in the Table 4. All RFU values of four GM maize genomes at all levels were higher than the corresponding threshold and revealed positive signal. The results suggested that four GM maize could be detected simultaneously at a starting template concentration as low as 0.5%, which was also lower than that in EU regulation and in consistent with singleplex analysis results. Therefore, the 4-plex MPFS method could be used to detect GM content with high sensitivity and high throughput.

**Table 4.**
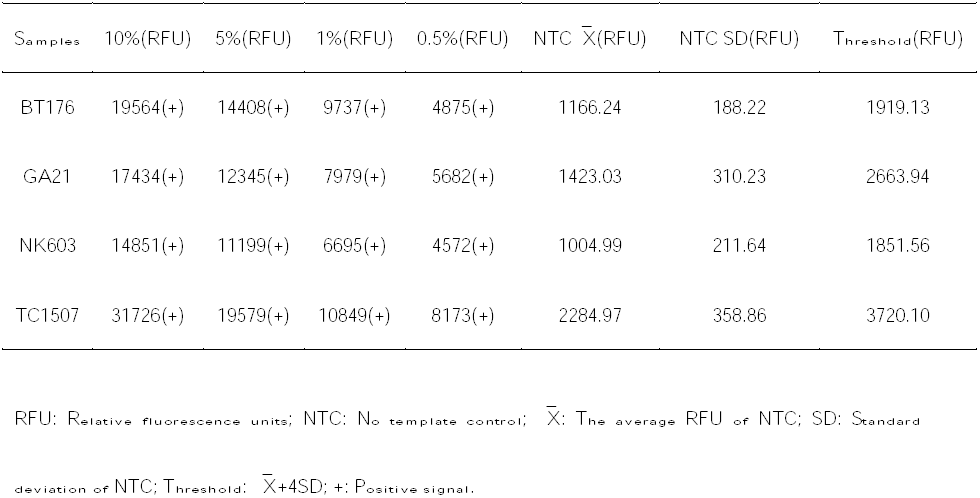
BT176, GA21, NK603 and TC1507 multiplex sensitivity assay.

#### Construction of Standard Curves and Quantitative Detection

Standard curves of 4-plex MPFS assay were established with a series of 5 dilutions (10^3^ – 10^7^copies) containing four GM maize. Results are shown in the Figure 2. All four GM maize showed an excellent linearity between the logarithm of RFU and copy numbers. The LOD of the 4-plex MPFS method is 10^3^copies. This is in good line with the singleplex assay, indicating the 4-plex MPFS PCR assay is of high accuracy and can be used to quantify the GM maize with high throughput.

**Figure 2.**
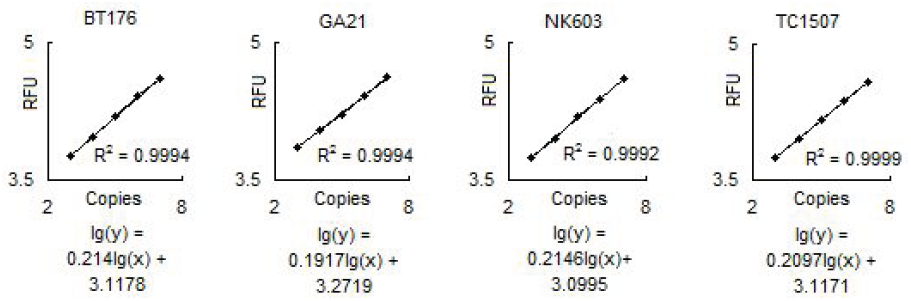
BT176, GA21, NK603 and TC1507 multiplex standard curves. RFU: Relative fluorescence units; Copies: The copies of plasmids.

### Repeatability

The repeatability of the 4-plex MPFS system for GM maize was measured by ten intra-assays and ten inter-assays at a copy number of 10^6^ in each assay. As the results shown in (Table 5), the coefficient of variation (CV) varied between 0.6-2.3% in the same batch and between 9.6-14.5% in the ten different batches, which fully satisfied the acceptance criterion.

**Table 5.** Repeatability of the RFU of four GM Maize standard plasmids at 10^6^ copies

| 106 Copies Plasmids | Intra-assay(RFU) |  |  | Inter-assay(RFU) |  |  |
| --- | --- | --- | --- | --- | --- | --- |
|  | AVE | SD | CV% | AVE | SD | CV% |
| BT176 | 39855.2 | 932.26 | 2.34 | 38027.7 | 3641.87 | 9.58 |
| GA21 | 38587.7 | 658.87 | 1.71 | 37467.2 | 4145.06 | 11.06 |
| NK603 | 38846.3 | 469.82 | 1.21 | 36877.9 | 145.06 | 11.24 |
| TC1507 | 38846.3 | 219.25 | 0.62 | 34649.3 | 5036.17 | 14.53 |

AVE: Mean relative fluorescence units; SD: Standard deviation; CV: Coefficient of variation; RFU: Relative fluorescence units.

### The Detection of Simulated Samples

To validate the accuracy and precision of MPFS method, four simulated samples were tested. Each sample was repeatedly amplified in triplex. After that, the mean RFU were determined and used to calculate the corresponding copies through the formula from the corresponding standard curve. The simulated percentage was calculated according to the ratio of the event-specific GM maize and the endogenous gene copy numbers. The GM contents calculated from experiments showed a slight deviation from the known GM contents (Table 6). Thus this method was accurate and precise enough to evaluate the GM contents in unknown samples.

**Table 6.** Quantification of four GM maize content in three simulated samples

| Samples | Fluorophores | RFU |  |  | AVE | SD | Copies | GM% |
| --- | --- | --- | --- | --- | --- | --- | --- | --- |
|  |  | 1 | 2 | 3 |  |  |  |  |
| BT176 MPFS assay |  |  |  |  |  |  |  |  |
| T1 (1%) | FAM | 16490 | 15282 | 15697 | 15823.00 | 501.15 | 113158.22 | 1.06 |
| T2 (3%) | FAM | 18842 | 19676 | 18961 | 19159.67 | 368.32 | 276694.01 | 2.95 |
| T3 (5%) | FAM | 22453 | 21604 | 21916 | 21991.00 | 350.64 | 526872.47 | 5.03 |
| GA21 MPFS assay |  |  |  |  |  |  |  |  |
| T1 (1%) | HEX | 16929 | 17966 | 17073 | 17322.67 | 458.69 | 110370.27 | 1.03 |
| T2 (3%) | HEX | 21560 | 19942 | 20470 | 20657.33 | 673.70 | 276504.32 | 2.95 |
| T3 (5%) | HEX | 22972 | 23029 | 23947 | 23316.00 | 446.79 | 519976.30 | 4.96 |
| NK603 MPFS assay |  |  |  |  |  |  |  |  |
| T1 (1%) | ROX | 14721 | 15764 | 14816 | 15100.33 | 470.88 | 107207.75 | 1.00 |
| T2 (3%) | ROX | 19064 | 18180 | 18214 | 18486.00 | 408.94 | 275183.45 | 2.93 |
| T3 (5%) | ROX | 20906 | 22019 | 20842 | 21255.67 | 540.39 | 527419.58 | 5.03 |
| TC1507 MPFS assay |  |  |  |  |  |  |  |  |
| T1 (1%) | NED | 15283 | 14942 | 14579 | 14934.67 | 287.45 | 109897.97 | 1.03 |
| T2 (3%) | NED | 17904 | 18415 | 18302 | 18207.00 | 219.16 | 282685.63 | 3.01 |
| T3 (5%) | NED | 20930 | 20956 | 20302 | 20729.33 | 302.36 | 524815.15 | 5.01 |
| IVR MPFS assay |  |  |  |  |  |  |  |  |
| T1 | CY5 | 39570 | 40607 | 40288 | 40155 | 433.67 | 10692504.1 |  |
| T2 | CY5 | 41284 | 36254 | 39960 | 39166 | 2128.86 | 9380519.15 |  |
| T3 | CY5 | 40822 | 40539 | 38636 | 39999 | 970.69 | 10476239.7 |  |

T1: 1% BT176, 1% GA21, 1% NK603, 1% TC1507, 96% Non-GM maize. T2: 3% BT176, 3% GA21, 3% NK603, 3% TC1507, 88% Non-GM maize. T3: 5% BT176, 5% GA21, 5% NK603, 5% TC1507, 80% Non-GM maize ;
RFU: Relative fluorescence units; AVE: Mean relative fluorescence units; SD: standard deviation; CV: coefficient of variation; GM% : Genetically modified maize content.

### Reproducibility

To evaluate the applicability of MPFS method, another four foodborne pathogenic bacteria were also tested. This 4-plex MPFS assay was performed and analyzed in the manner of MPFS system described above except for the PCR amplification system.

#### Specificity Assay, Qualitative Detection and High Throughput

The DNA mixtures containing four foodborne pathogenic bacteria were used as template to evaluate the specificity of the 4-plex MPFS again. Results showed that the RFU values from four foodborne pathogenic bacteria genome amplifications were higher than the corresponding threshold (Table 7). Therefore, the 4-plex MPFS PCR assay for four foodborne pathogenic bacteria showed the same specificity as the MPFS assay for four GM maize. This indicates that the MFPS method can also be applied to qualitatively detect multiple genes in other biological samples.

**Table 7.** Multiplex foodborne pathogenic bacteria specificity assay.

| Samples | Genome(RFU) | NTC $\bar{X}$(RFU) | NTC SD(RFU) | Threshold |
| --- | --- | --- | --- | --- |
| Salmonella | 33994(+) | 1316.52 | 222.11 | 2204.96 |
| L. monocytogenes | 12538(+) | 1826.74 | 235.13 | 2767.25 |
| E. coli | 22447(+) | 882.10 | 175.11 | 1582.53 |
| S. aureus | 13278(+) | 3026.25 | 562.51 | 5276.27 |

RFU: Relative fluorescence units; NTC: No template control; $\bar{X}$ : The average RFU of NTC; SD: Standard deviation; Threshold: $\bar{X}+4SD$ ; +: Positive signal; L. monocytogenes: Listeria monocytogenes; E. coli: Escherichia coli; S. aureus: Staphylococcus aureus.

#### Construction of Standard Curves and Quantitative Detection

Standard curves of 4-plex MPFS PCR assay for four foodborne pathogenic bacteria were established again with mixed four DNA dilutions. The logarithm of RFU values and the target gene copy number showed excellent linear relationship for all four bacteria (Fig. 3). The LOD was determined to be 10^3^ copies. Again, the results are in good line with the results of 4-plexplex MPFS assay for four GM maize samples, implying its potential use in quantifying the genes in other biological samples with high throughput.

**Figure 3.**
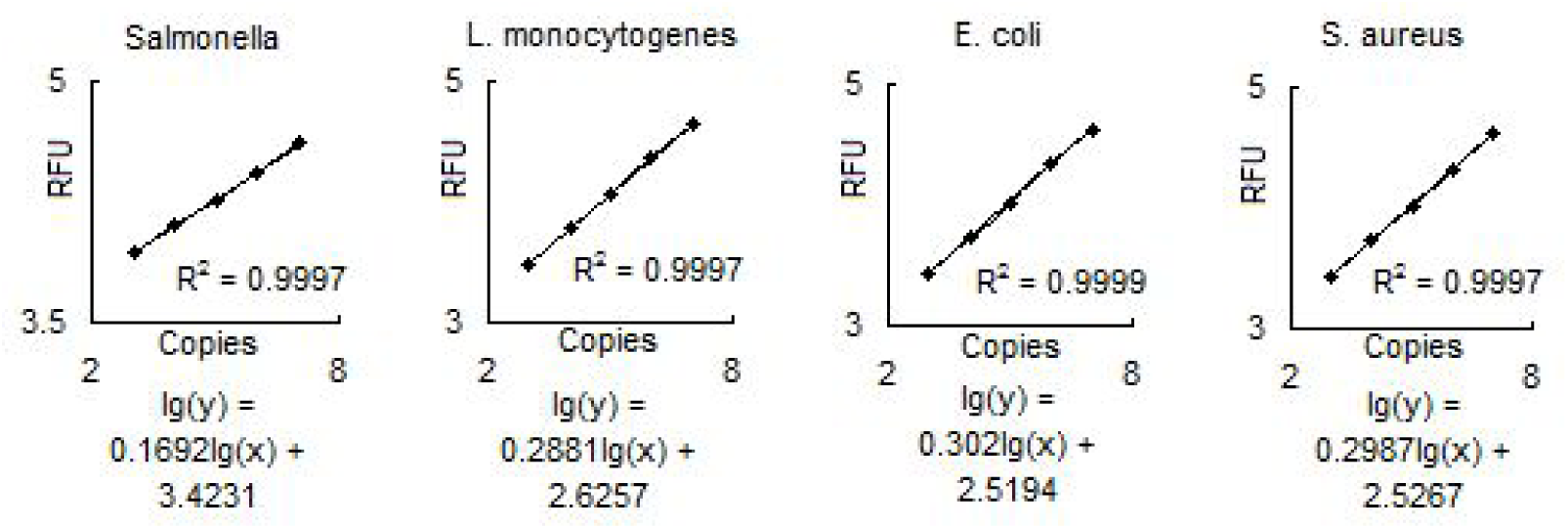
four foodborne pathogenic bacteria multiplex standard curves. RFU: Relative fluorescence units; Copies: The copies of plasmids; L. monocytogenes: Listeria monocytogenes; E. coli: Escherichia coli; S. aureus: Staphylococcus aureus.

#### Repeatability

The repeatability of the MPFS system for four foodborne pathogenic bacteria were studied by ten intra-assays and ten inter-assays for four foodborne pathogenic bacteria at a copy number of 10^6^. The coefficient of variation (CV) was determined between 1.2-2.2% in the same batch and between 9.1-11.8% among the ten different batches, in good line with the repeatability of the four GM maize assay (Table 8).

**Table 8.** Repeatability of the RFU of four foodborne pathogenic bacteria standard plasmids at 10^6^ copies.

| 10<sup>6</sup> Copies Plasmids | Intra-assay |  |  | Inter-assay |  |  |
| --- | --- | --- | --- | --- | --- | --- |
|  | AVE | SD | CV% | AVE | SD | CV% |
| Salmonella | 36065 | 963.43 | 2.67 | 36159.8 | 4253.83 | 11.76 |
| L. monocytogenes | 39208.3 | 579.29 | 1.48 | 37504.1 | 3400.74 | 9.07 |
| E. coli | 38827.9 | 473.30 | 1.22 | 39647.6 | 4231.33 | 10.67 |
| S. aureus | 38952.9 | 596.13 | 1.53 | 35523.1 | 4147.47 | 11.68 |

AVE: mean relative fluorescence units; SD: standard deviation; CV: coefficient of variation;

Altogether, the results demonstrated good specificity, sensitivity, linearity and repeatability of this MPFS method developed in our laboratory, which could be used for qualitative, quantitative and high-throughput detection and analysis of multiple genes in a single reaction.

## Discussion

In this study, we combined microdroplet PCR with fluorescence spectrophotometry (MPFS) and applied it to qualitative, quantitative and high-throughput detection of four target DNAs. GM maize was selected as a model for this purpose. By means of labeling the different gene-specific primer with different fluorophores without detection interference, multiple target DNAs could be co-amplified in a single PCR reaction, and amplicons could be analyzed qualitatively and quantitatively with high throughput. The method was proved to be free of cross interference from different fluorophores, and of sensitivity of 0.5% (w / w) GM content, lower than 0.9% of EU criterion. In addition, this method generates the data with excellent linearity relationship between the logarithm of RFU values and gene copy number and with a LOD of 10^3^ gene copy number. The 4-plex method increased the throughput by microdroplet PCR reaction from single MPFS assay but not sacrifice the quality of resulting data. The small signal strength variation within batch or between batches further confirmed the precision and accuracy of this MPFS method. Finally, the results from foodborne pathogenic bacteria showed the applicability of MPFS method to other biological samples.

The excellence of MPFS method is apparently attributable to the combination of emulsion PCR and fluorescence spectrophotometry. Microdroplet PCR allows for improving the throughput, which avoids constraints in multiple PCR by partitioning the reaction mixture into discrete droplets while remaining the specificity and sensitivity. Microdroplet PCR, also called emulsion PCR, was first systematically described by Williams et al. to amplify complex DNA mixtures by compartmentalization of genes in water-in oil (w/o) emulsion (Williams et al. 2006). Emulsion PCR method was consisted of emulsion PCR and analysis of the products. Williams et al. used agarose gel electrophoresis to analyze the PCR products. But Williams’ method can only be used for qualitative assay and not for quantitative assay (Williams et al. 2006). And the contamination derived from the agarose gel electrophoresis was a defect. Guo et al also used microdreoplet PCR implemented capillary gel electroresis to amplified and analyze multiple DNA targets (Guo et al. 2011). However, the two-step method is relative complicated and it can not used for quantitative assay too. Droplet digital PCR enables absolute quantitation of low copies sample, but it needs skilled operators and expensive consumption and the results is astable (Hindson et al. 2011). However, we combined microdroplet PCR with fluorescence spectrophotometry to amplify and analyze multiple targeted DNA qualitatively and quantitatively with high throughput. Conventional PCR could only be used for qualitative detection, but can not be used for quantitative detection and the throughput was limited (Germini et al. 2004). In the MPFS method, the PCR products were directly measured by Infinite M1000 PRO to avoid the contamination derived from the agarose gel electrophoresis. Both the real-time quantitative PCR and our MPFS method employed fluorescence labeling technique. Real-time quantitative PCR enables quantitative detection with low LOD and high accuracy, it has, however, limit in the throughput due to limited detecting channels (Chaouachi et al. 2013). The MPFS method used Infinite M1000 PRO to measure the RFU of the fluorescence, which allows for detecting emission wavelength from 200 to 1000nm (Thermo VarioSkan Flash), that is, 12 fluorophores can be analyzed in a well. With appropriate adjustment, almost 20 fluorohpores could be tested simultaneously in a well, which means more than 6000 genes can be detected in a 384-well plate at a time. Therefore, our method could detect abundant target genes simultaneously. DNA microarray is good for high-throughput and qualitative analysis, but not good for quantitative assay. The DNA microarray also uses fluorescence labeling technique, however, the problems in biochip microfabrication technology limits its popularization and application. Also, preparation of high-density array was time consuming and the confocal laser scanner is expensive (Hessner et al. 2003). In the process of amplification with microarray, the target sample was found easy to be contaminated and the signal-to-noise would be affected (Hessner et al. 2003). As a contrast, MPFS method needs only conventional PCR thermocycle instrument and a fluoro-microplate reader (or fluorospectrophotometer) to detect multiple target DNAs qualitatively and quantitatively with high throughput, clearly indicating its association with simplicity, speediness, concision and low cost.

Emulsion PCR were performed according to the method of Richard et al. (Williams et al. 2006) with some improvement such as the time interval every two drop (6s), the composition of the oil phase, the 3 × 8 mm stir bar, the ampoule with a lid and the adoption of centrifugation to break the emulsion without other process.

Many emulsion PCR techniques currently adopted combine with a known instrumentation and technologies for DNA analysis, such as agarose gel electrophoresis (Williams et al. 2006), microarray (Ge et al. 2006), capillary electrophoresis (Guo et al. 2011), micro-fluidic chip (Zhu et al. 2012). Our new analysis method for multiple target DNAs, fluorescence spectrophotometry, has been not reported previously. The combination of fluorescence spectrophotometry with the emulsion PCR, to our knowledge, is the first report to detect multiple target DNAs qualitatively and quantitatively with high throughput. The excellent performance of the MPFS method enables its application in the fields of animal husbandry and veterinary pathogenic microorganisms, human pathogenic microorganisms, aquatic pathogenic microorganisms, environmental microbes and plant pathogenic microbes. The MPFS method could also be used for detection of varieties of human disease-related genes and genetic counseling.

The brilliant performance of MPFS system described in this study is still dimmed by the manual purification of emulsion products. This, however, can be improved by the use of automatic purification system, by then the repeatability and accuracy will be greatly improved.

## Materials and Methods

### Materials

GM maize flour i.e., Bt176, GA21, NK603 and TC1507, were supplied by Monsanto Company (St. Louis, MO). Non-transgenic seeds (maize soybean, rapeseed, cotton, wheat) were purchased from a local market in Shanghai, China. Four common foodborne pathogenic bacteria i.e., Salmonella, Listeria monocytogenes, Escherichia coli and Staphylococcus aureus DNA and plasmids used for reproducibility assay were obtained from this laboratory.

### Sample Preparation

The genomic DNA of all plant materials were extracted and purified using a mini-plant genomic DNA extraction kit (Shanghai Bio-ful Biotech Co., Ltd., Shanghai, China). The DNA concentration and quality were estimated using a NanoDrop 1000 UV-Vis spectrophotometer (NanoDrop Technologies, LLC, Wilmington, DE).

To test the sensitivity, a series of targeted GM DNA solution were obtained by mixing the GM maize flour with non-GM maize flour. The final relative GM maize content was 10%, 5%, 1%, 0.5% (w / w) for four GM maize, respectively. Moreover, event-specific genes were used for the construction of standard plasmids as calibrators for GMO quantification and represented an alternative to genomic DNA. Additionally, various GM contents (1%, 3% and 5% GM, w / w) of four simulated samples were also prepared to evaluate the accuracy and precision of MPFS system. Easy dilution (TaKaRa biotechnology Co., Ltd, Dalian, China) was used to dilute the standard plasmids to avoid DNA loss due to adsorption to the tube walls.

### Set-up of MPSF Reaction

#### The selection of Fluorescence

Fluorescence labeling technique was used to label primers. When the sum of excitation and emission bandwidth was less than stokes shift of a fluorophore, the optimal excitation and emission wavelength could be set to get stronger signal and prevent overlap. Based on parameter characteristics of the Tecan Infinite M1000 PRO and Thermo VarioSkan Flash, the excitation bandwidth was 5nm and the emission bandwidth was 12 nm. So the fluorophores, whose stokes shift were more than 17nm, were more suitable to be selected. Furthermore, the excitation spectrum and emission spectrum of two adjacent fluorophores should not be overlapped either. On the basis of the above principles, twelve fluorophores were selected from more than 300 fluorophores which can be used at the same time. The excitation and emission wavelength were properly adjusted without affecting the fluorescence strength and sensitivity, then almost 20 fluorohpores could be tested simultaneously. In this study, four fluorophores were elaborately selected and four pairs of primers were labeled with selected fluorophores (Table 9) and synthesized (Thermo Fisher Scientific Co., Ltd., Shanghai, China) to amplify the targeted DNA.

**Table 9.**
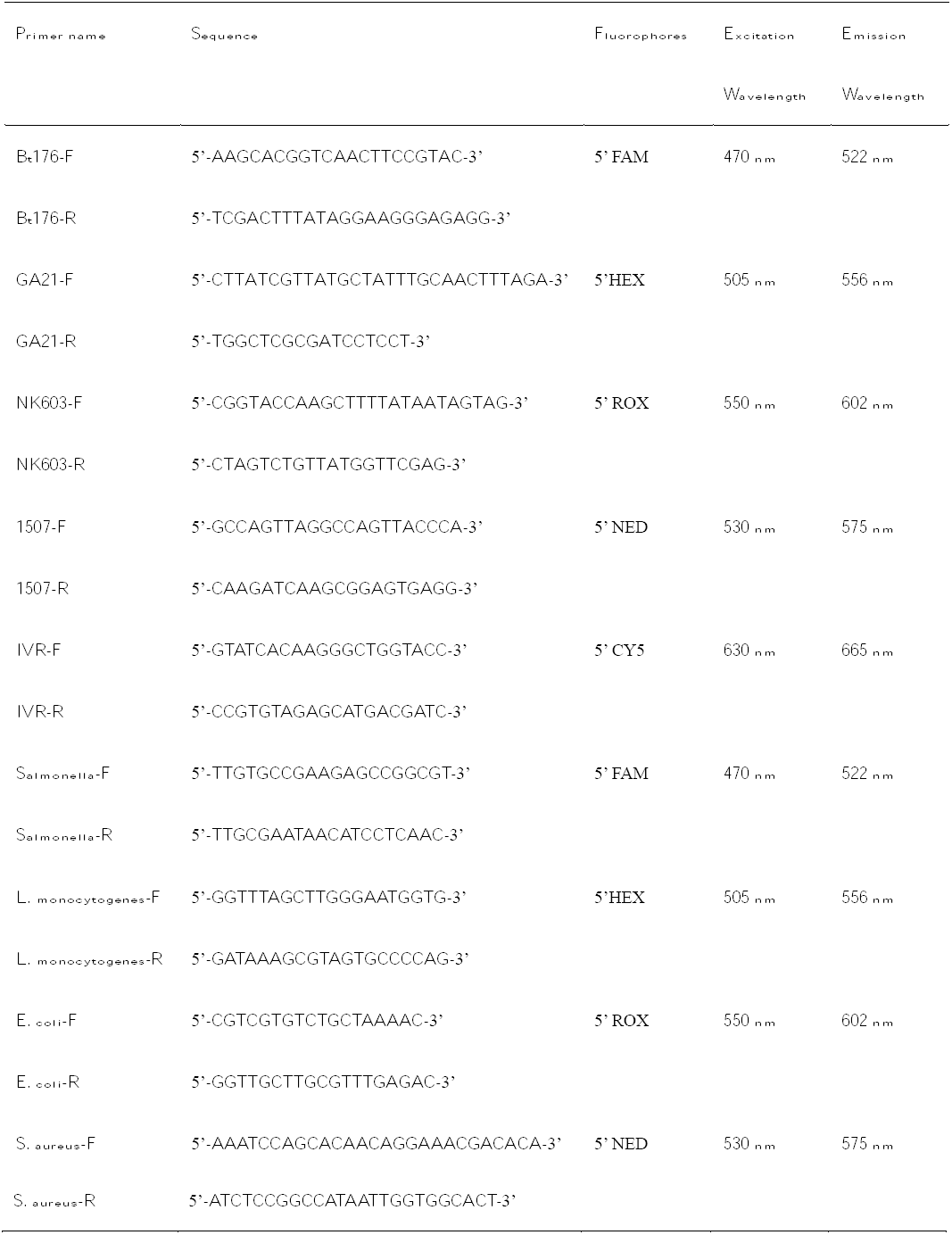
Primer sequences and excitation and emission wavelength of fluorophores.

#### Singleplex PCR Composition

The aqueous phase for four GM maize in individual reaction consisted of 1 × PCR buffer, 10g/L BSA, 0.2mM dNTPs, 0.4 uM of forward and reverse primers, 26U Taq DNA polymerase and 10.4 μL template DNA in a total volume of 260uL.

#### Multiplex PCR Composition

Four single PCR for four GM maize were pooled into one reaction with the same composition except for the amount of primers and Taq polymerase. The optimized primers and templates were as follows: 0.2uM for each pair of forward and reverse primers and 104U for Taq polymerase.

Multiplex PCR aqueous phase for four foodborne pathogenic bacteria consisted of 10×PCR buffer, 10g/L BSA, 0.2 mM Mg^2+^, 0.3 mM dNTPs, 0.2 uM for each pair of forward and reverse primers, 16.25U Taq DNA polymerase and 6.5 μL template DNA in a total volume of 260uL.

#### Microdroplet PCR Amplification

The emulsion PCR was carried out according to a previously described emulsion PCR-based method (Williams et al. 2006) with several modifications. The oil-surfactant mixture was prepared by mixing the 4.5ml span 80, 0.4ml tween 80, 0.05ml triton X-100 and 95.05 ml mineral oil at 1000 r.p.m for 2h. At 1000r.p.m, the water – in-oil emulsion were obtained through adding 200ul of the aqueous phase to 400ul oil phase drop-wise (time interval is 6s) while stirring with a magnetic stirring bar (3 × 8 mm) in a disposal 2ml ampoule with a lid at 25 °C. Over the period of 5min, the water – in-oil mixture is processed into discrete encapsulation of individual reaction. After that, the obtained w/o emulsion was delivered into PCR tubes as 12 aliquots of 50 μL for further thermal cycling and residual 50 ul of the aqueous phase were used as a no emulsified control. The PCR for four GM maize were performed as the following program: 95°C for 5min; 35 cycles at 94°C for 30s, 57°C for 40s, 68°C for 35s; a final extension at 68°C for 7min and then hold at 4°C. The PCR protocols for four foodborne pathogenic bacteria were as the follows: 94°C for 5min; 35 cycles at 94°C for 30s, 60°C for 30s, 72°C for 30s; a final extension at 72°C for 7min and then hold at 4°C

Prior to the standard analyzing protocol for the PCR products, the resulting emulsion mixtures were pooled and centrifuged at 16200g for 5min. Approximately 140ul of lower aqueous phase contained fluorescent target DNA were then obtained and purified to remove the residual primers. An AxyPrep PCR Cleanup Kit (Axygen Scientific, Inc.) was used in accordance with the manufacturer’ s instructions except for eluting with 100 μL elution buffer to get the final amplicons for subsequent analysis.

### Infinite 200^®^ PRO and Magellan analysis

Magellan Standard software and Infinite M1000 PRO (Tecan Austria GmbH, Grödig, Austria) possessing narrow bandwidths were used for multiple labeling analyses in line with the manufacturer’ s standard instructions. The resulting samples from PCR were directly transferred to a disposal, black, flat-bottom, 384-well Fluorotrac^TM^ 200 plates (Greiner Bio-One Suns Co., Ltd, Beijing, China). Fluorescent intensity scan were performed to determine the optimal exitation and emission wavelength (Table 5). Top-read determination of fluorescence intensity was performed for qualitative, quantitative and high-throughput analysis. Due to the intrinsic excitation and emission wavelength of fluorescence, the RFU of four kinds of targets DNA harboring four fluorophores could be measured respectively. The manual gain was set to 100. All the data generated was derived to Excel to conducted relative analysis.

#### Fluorescence Intensity Determination

In Bt176’ s case, fluorescence intensity was performed on the resulting DNA samples to be analyzed with the flowing parameters: excitation wavelength λ =470nm and emission wavelength λ =522nm. A gradient of standard samples and the mapping concentration were set to construct the standard curve. Furthermore, the linear regression analysis, the correlation coefficient and blank reduction were adopted to provide a net RFU increase which would be proportional to DNA copies in the resulting products.

### Calculation of threshold

Following the MPFS method we developed, 1000 negative samples were used for determination of the threshold. Based on statistically analysis of the RFU from negative, they form a normal distribution.

As a threshold signal value, a level mapping to blank (no template control, NTC) 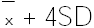 was chosen as criterion for a positive signal. The probability density of the normal distribution is:

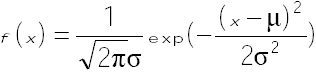

Where μ is the mean or expectation of the distribution and also its median and mode 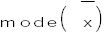, σ is the standard deviation (SD), σ^2^ is the variance, and x is the independent variable for which you want to evaluate the function.

## Acknowledgments

This work was financially supported by Shanghai Minhang Science Committee foundation of China, Major projects of industrialization, grant no. 2014MH076.

## Author contributions

K.Z. conceived the initial hypothesis, designed experiments, wrote and edited the manuscript, K.Z., Y.N.D., X.Z., W.S. and B.A.Z. performed most of the experiments, F.F.R, Y.Y.W. and R.L.H. performed some of the experiments, B.S. and H.Z.Z contributed reagents/materials/analysis tools, Y.X., X.R.F., Q.Z. and X.X.Z. analyzed the data, Y.N.D. and X.Z. wrote the manuscript.

## Conflict of interest

The authors declare no competing interests.

